# Development and pilot application of a point-of-need molecular xenomonitoring protocol for tsetse (*Glossina sp.*) in a low-resource setting

**DOI:** 10.1101/2025.11.10.685795

**Authors:** Isabel Saldanha, Edward Aziku, Alex T. Trima, Victor Drapari, Gala Garrod, Henry Ombanya, Inaki Tirados, Martha Betson, Albert Mugenyi, Sophie Dunkley, Stephen J. Torr, Andrew Hope, Lucas J. Cunningham

## Abstract

**Background:** Tsetse flies (*Glossina sp.*) are the primary vectors of trypanosomes causing human African trypanosomiasis (HAT) and animal African trypanosomiasis (AAT). Disease surveillance can be carried out by detecting *Trypanosoma* DNA in tsetse, also known as molecular xenomonitoring. Whilst molecular methods can increase the efficiency and sensitivity of pathogen detection, trained staff and a well-equipped laboratory are required. In many cases, DNA extraction and screening is outsourced to a central laboratory in a major city either in-country or abroad, far removed from original tsetse collection sites. This increases results turnaround time, incurs transportation costs, and can lead to sample loss or damage.

**Methodology/Principle Findings:** We set out to develop, optimise and trial methods for tsetse xenomonitoring in a low-resource point-of-need setting. A low-cost protocol was developed consisting of rapid alkali-based DNA extraction and *Trypanosoma* detection qPCR assays using air-dryable reagent mixes. A minimally-equipped laboratory was established in a field station in Arua, Uganda. Following a training workshop, three entomology technicians carried out screening on 286 tsetse collected over a nine-week study period. The technicians consistently extracted high quality DNA (98% success rate) and were able to successfully detect *T. brucei sensu lato*, *T. congolense* and *T. vivax* DNA in 3.6% - 4.3% (95% confidence interval [1.73, 7.73]) of total tsetse.

**Conclusions/Significance:** This study demonstrated that sensitive molecular xenomonitoring of HAT and AAT pathogens can be carried out without the need for cold-chain storage or high-powered equipment. Further improvements to the system might be achieved by modifying the DNA extraction protocol to enable high-throughput or pooled samples, increasing the sensitivity of the *T. b. gambiense* DNA detection assay and exploring more sustainable power sources.

**Author Summary:** Tsetse flies spread the parasitic diseases human African trypanosomiasis (sleeping sickness) and animal African trypanosomiasis (nagana) that impact populations across sub-Saharan Africa. Disease surveillance can be carried out using tests to detect parasite DNA in tsetse, termed molecular xenomonitoring. Currently, these methods are too complex, costly and logistically-challenging to be carried out in remote areas where sleeping sickness is a problem. However, advances in molecular testing technology are now making this a possibility. We set out to develop a tsetse molecular xenomonitoring system using a basic laboratory set-up in Arua, Uganda. The protocol comprised a low-cost method to extract DNA from tsetse, a portable qPCR machine to test samples and air-dried reagents that did not require cold storage. Following a two-week training workshop, three technicians went on to carry out testing on 286 tsetse over a nine-week period. The technicians were able to consistently extract high-quality DNA (98% success rate) and successfully detected trypanosome parasite DNA in 30 (10.7%) tsetse samples. Whilst there are still challenges to overcome, this study has demonstrated that molecular xenomonitoring of tsetse can be carried out without the need for trainees with previous molecular experience, refrigerated reagents or high-powered equipment.

## Introduction

Tsetse flies (*Glossina sp.*) are the primary vectors for trypanosomes that cause the neglected tropical disease human African trypanosomiasis (HAT) and the veterinary disease animal African trypanosomiasis (AAT). HAT is caused by two subspecies of *Trypanosoma brucei* (of the sub-genera *Trypanozoon*); *T. b. rhodesiense* is the cause of East African ‘Rhodesian’ HAT (rHAT) and *T. b. gambiense* causes West African ‘Gambian’ HAT (gHAT). AAT is caused by three different species of trypanosome; *T. b. brucei* (of *Trypanozoon*), *T. congolense* (of *Nannomonas*) and *T. vivax* (of *Duttonella*).

Uganda lies squarely within the 10 million km^2^ of sub-Saharan Africa infested by tsetse. The dominant vector species is *G. fuscipes fuscipes* found in riverine habitats, whilst *G. morsistans morsitans* and *G. pallidipes* are found in savannah habitats [1]. Uniquely, Uganda is the only country to have had known foci of both forms of HAT; rHAT in south-eastern districts, and gHAT in the north-west [2] across nine districts: Adjumani, Amuru, Arua, Koboko, Maracha, Moyo, Nebbi, Yumbe and Zombo. Once the cause of devastating epidemics resulting in an estimated 300,000 to 500,000 deaths at the turn of the 20^th^ century [3], gHAT has now been eliminated as a public health problem in Uganda [4] in line with WHO 2030 elimination targets [5]. AAT however remains a significant animal health and economic burden across sub-Saharan Africa, including Uganda [6]. Livestock surveys in the Uganda’s West Nile region have identified AAT aetiological agents in cattle, pigs, sheep, goats and dogs [6–9] in addition to local tsetse populations [7,9–13]. Livestock ownership and production is rising across Uganda. According to the 2021 Uganda National Livestock Census, the total cattle population grew by 27% to 14.5 million animals from 2008 to 2021 [14], whilst goat, sheep and pig populations grew by 39%, 28% and 13% respectively during the same period [14]. At the same time, AAT prevalence estimates are limited at district-level in Uganda [6]. Therefore, there is a need for improved and timely AAT surveillance across all districts.

Pathogen surveillance in vectors plays a critical role in a disease elimination or post-elimination setting to confirm the absence of transmission [7,15–18]. Systematic sampling of tsetse populations allows not only the monitoring of tsetse population dynamics but is also a way of estimating parasite prevalence in a particular area. Historically, individual tsetse were collected, dissected and subjected to microscopic analysis to determine whether *Trypanosoma sp.* were present, with species identification dependent on which vector tissues were colonised [19]. This technique was the gold standard for identification of trypanosome infection in tsetse for several decades and is still in use today [19–21]. Classic microscopic detection is increasingly being replaced by detection of parasite DNA, known as ‘molecular xenomonitoring’. Microscopy is labour-intensive and suffers from poor sensitivity and specificity due to limitations in microscope resolution and similarities in *Trypanosoma* morphology [20–23]. Molecular xenomonitoring has the advantages of much higher diagnostic sensitivity and specificity and the potential for high-throughput sample analyses [13,20]. These are particularly beneficial traits for HAT and AAT xenomonitoring, where parasite incidence in tsetse populations can be as low as 0.1% [20,24], necessitating large sample sizes.

There is growing demand for molecular diagnostics and surveillance at the point-of-need (PON) for many neglected tropical diseases [25]. In the case of xenomonitoring, samples are usually processed and analysed in a centralised laboratory, often far removed from the original collection sites. This can lead to sample loss or damage during transportation, increases the time-to-result and can restrict knowledge transfer and capacity building at the PON. In the case of shipments to other countries, samples may also be subject to export/import restrictions and to the Nagoya Protocol [26]. Despite a clear need for building molecular diagnostic capacity, HAT and AAT molecular diagnostics and xenomonitoring at the PON still has many well-documented challenges [27–29]. Intermittent power supply, lack of resources, variable user knowledge and experience, uncontrolled temperature and humidity and presence of possible contaminants or inhibitors are all potential barriers to implementation. The World Health Organisation target product profile (TPP) for a gHAT diagnostic test states a need for minimal reliance on a cold chain, specimen preparation involving the least possible precision liquid handling and laboratory training that can be delivered within a week [27]. These targets should also be kept in mind when designing PON xenomonitoring protocols.

Tsetse xenomonitoring presents its own additional challenges. DNA extraction is a major hurdle across all PON diagnostics [30] and the choice of method can impact DNA yield, quality and downstream analyses [31]. Whilst rapid DNA extraction protocols for liquid samples are well-established [30], tsetse tissue normally requires mechanical lysis, heating and/or overnight incubation steps [7,11,12,24]. Trialled ‘field-friendly’ DNA extraction methods for arthropods have ranged from manual crushing in sterile distilled water [32], to use of FTA cards [33,34], lysis buffers [35], alkaline extraction [36] and commercial reagents or buffer-based kits [37,38]. Nevertheless, these methods have not yet been trialled in larger arthropods such as tsetse.

A myriad of molecular assays have been developed to detect *Trypanosoma sp.* DNA across PCR, qPCR and LAMP platforms [20], reflecting the complex nature of identifying *Trypanosoma sp.* in tsetse and their hosts. Assays that have been used to detect *T. b. gambiense* DNA in tsetse include PCR, LAMP and HRM [11,39–41]. The greater species-diversity of AAT requires a PCR or restriction fragment length polymorphism (RFLP) that targets the conserved internal transcribed spacer (ITS) region [42–44], or three (or more) species-specific PCR or qPCR assays [44–46]. The diverse range of non-target *Trypanosoma sp.* and related flagellates that can be detected in tsetse, coupled with the possibility of co-infections of multiple trypanosome species [47,48] can reduce diagnostic specificity. Therefore, the choice of xenomonitoring assay(s) and platform must be carefully considered.

Accordingly, we set out to design, optimise and test a molecular xenomonitoring protocol in a low-resource point-of-need setting. Key components would include a low-cost alkali and magnetic bead-based DNA extraction and purification method [10,49], an air-dryable qPCR mix (PCR Biosystems Ltd, London, UK) that did not require cold-chain and a small and highly portable qPCR machine (Mic, Bio Molecular Systems, Upper Coomera, Australia). Our principal objectives were the production of a robust, sensitive and easy-to-use protocol to detect *T. b. gambiense* (gHAT), *T. b. rhodesiense* (rHAT) and *T. b. brucei*, *T. congolense* and *T. vivax* (AAT) in a wild tsetse population over several weeks.

## Methods

The project was carried out in two stages: a protocol development phase carried out in the UK, followed by a trial phase carried out in Uganda (Figure 1).

**Figure 1:**
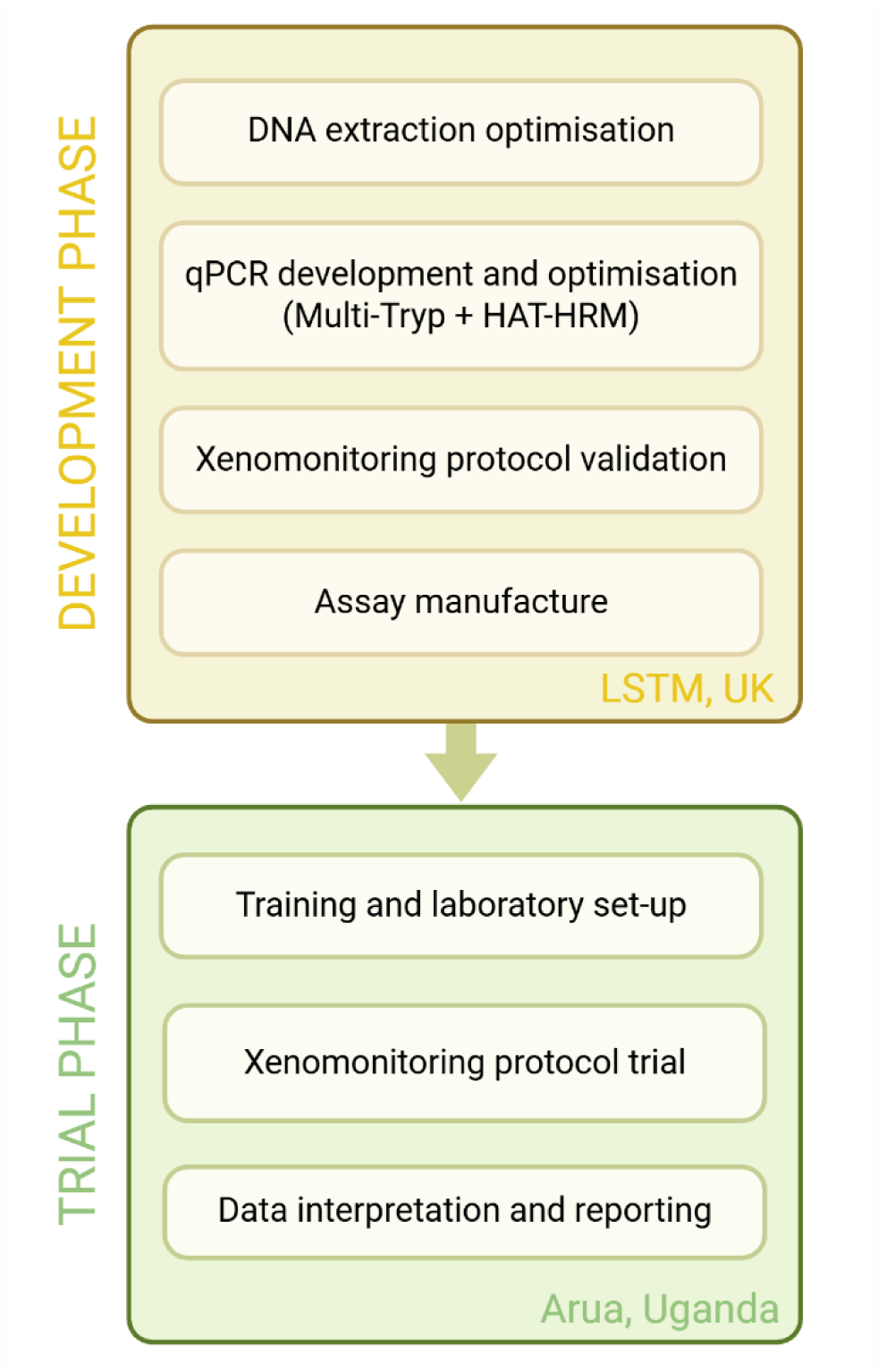
A flow diagram illustrating the protocol development and trial process. Image created using biorender.com [accessed 23/04/25].

### Protocol development and optimisation

#### Field DNA extraction optimisation

To assess potential field DNA extraction methods, three test methods (TM1-3) were compared against a ‘gold standard’ method (QIAGEN DNeasy Blood and Tissue kit, QIAGEN, Germany) detailed below. Test methods were as follows: TE buffer (TM1), lysis buffer (TM2) and alkaline (S1 Text) extraction (TM3). Each test method included a purification step using magnetic beads (SpeedBead Mix; S1 Text) developed by Byrne *et al* [10,49]. For each test method and gold standard, nine female insectary-reared *G. m. morsitans* (8-9 weeks old) were dissected, 24 hours post-bloodmeal, and processed following the protocols outlined in S1 Figure.

#### ‘Gold standard’ DNA extraction

For insectary-reared flies, whole intact tsetse or total dissected remains were placed into individual collection tubes and incubated at 56°C for 3 hours on a rocker set at 5 oscillations per minute to remove ethanol. DNA was extracted and purified using QIAGEN DNeasy 96 Blood and Tissue Kit (QIAGEN, Hilden, Germany) following the manufacturer’s protocol, slightly optimised for large insect processing by the addition of a mechanical lysis step. In short, after ethanol removal, a 6.35 mm diameter stainless-steel ball (Dejay Distribution Ltd, UK) was placed into each tube. After adding Buffer-ATL/Proteinase K, samples were then mechanically lysed at 15Hz for 20 seconds for two rounds using a QIAGEN TissueLyser II (QIAGEN, Hilden, Germany). Following centrifugation at 2000rpm for 1 minute, samples were incubated at 56°C for 14 hours. Eventual purified DNA was eluted in 80µL elution buffer AE.

#### Primer and probe design

For TCF-qPCR, two *T. congolense* Forest minichromosomal satellite DNA entries (accession numbers S50876.1 and JX910380.1) were retrieved from GenBank. Multiple sequence alignment was performed using Clustal Omega and analysed using Jalview v2. The resulting consensus sequence was used as the template for primer and probe design. For TVX-qPCR, the *T vivax* minichromosomal satellite DNA entry (accession number J03989.1) was used as a template for primer and probe design. For UGWigg-qPCR, a previously-sequenced *Wigglesworthia glossinidia* partial 16s rRNA gene amplicon (GenBank accession number PV800902.1) obtained from an adult uninfected *G. f. fuscipes* individual collected from West Nile region [50] was used as a template for primer and probe design. All qPCR primers and probes were designed using Primer3 and *in silico* primer sequence specificity analysis was performed using BLAST (NCBI). Primer and probe sequences are in Table 1.

**Table 1:**
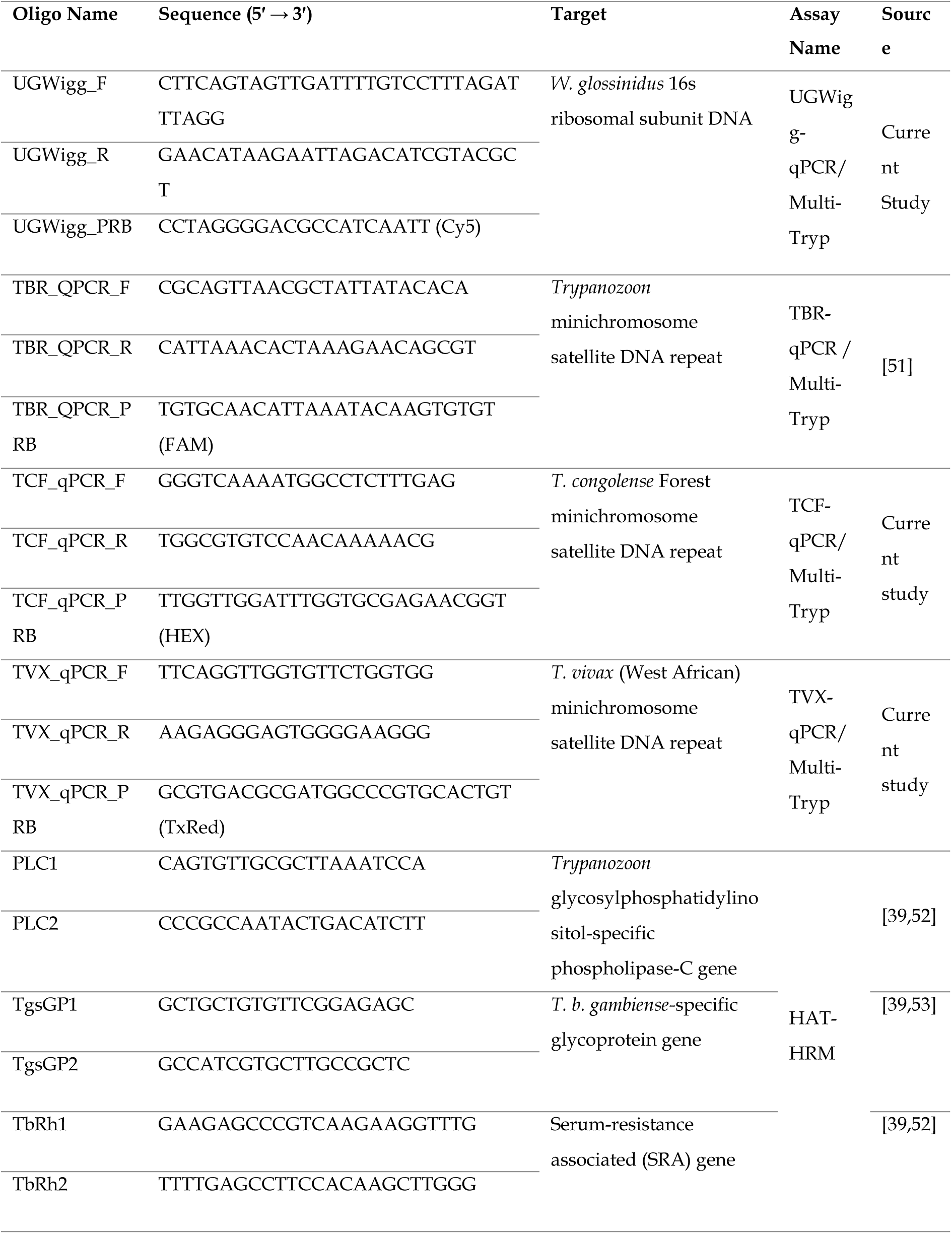
Primers and probes used in the study.

### qPCR optimisation and validation

All qPCR reactions were carried out using PCR Biosystems Air-Dryable Probe Mix (PCR Biosystems Ltd, London, UK) in line with the manufacturer’s protocol. All primer and probe sequences are shown in Table 1.

During optimisation, a ‘wet mix’ was used consisting of 5 µL template DNA mixed with 2.5 µL of 4X Air-Dryable Probe Mix, 0.2 - 0.6 µM forward and reverse primers, 0.1 – 0.2 µM probe (Multi-Tryp) or 0.5 µL EVAgreen (HAT-HRM) and 0.3M betaine (HAT-HRM) and nuclease-free water (NFW) added to a 10 µL total reaction volume.

The ‘dry mix’ consisted of 2.5 µL of 4X Air-Dryable Probe Mix (PCR Biosystems Ltd, London, UK) added to 0.2 - 0.6 µM forward and reverse primers, 150-200 µM probe (Multi-Tryp) or 0.5 µL EVAgreen (HAT-HRM) and 0.3M betaine (HAT-HRM) resulting in a 3.88 µL (Multi-Tryp) and 4.80 µL (HAT-HRM) total reaction volumes. After the master mix was complete, 3.88 µL (Multi-Tryp) and 4.80 µL (HAT-HRM) of these solutions were then aliquoted into each upturned qPCR tube lid. Lids were then dried at 40°C for 80 minutes in a hybridisation oven (Cole-Parmer Stuart, US) before packaging in airtight sealable polythene bags containing ∼5g colour-indicating silica gel (Fisher Scientific, Hampton, US), labelled with assay name and wrapped in UV-protective aluminium foil.

When required, the dry mix was rehydrated by adding 5 µL of NFW to each qPCR tube cap containing desired assay dry mix and left to incubate within a dust-protective box at room temperature (RT) for 10 minutes. In the interim, 5 µL of each extracted DNA sample (or 5 µL NFW for negative template control; NTC) was added to the corresponding qPCR tube(s). Tubes were then carefully capped and liquids mixed by flicking tubes and inverting tube-cap assembly three times.

Thermal cycling conditions were as follows; initial denaturation at 95°C for 3 minutes followed by 40 cycles of denaturation at 95°C for 15 s and annealing and extension at 58 - 61°C for 35 s (Multi-Tryp) or 58°C for 35 s (HAT-HRM), followed by a high-resolution melting step between 65 – 95°C at a rate of 0.1°C (HAT-HRM only). Data were captured during the annealing and extension step. Thermocycling, fluorescence detection and data capture was carried out using a Mic and micPCR v.2.9.0 software (Bio Molecular Systems, Upper Coomera, Australia).

Analytical sensitivity and specificity experiments were carried out for all assays in wet and dried formats. All optimisation and validation experiments were carried out in triplicate. A composite DNA sample, consisting of previously-extracted DNA from 284 wild uninfected *G. f. fuscipes* sourced from Maracha District [10] was used as a positive control for *G. f. fuscipes*-specific *W. glossinidia* DNA. Analytical specificity testing was carried out against a panel of nine *Trypanosoma sp.* DNA samples and the *G. f. fuscipes* composite DNA sample.

A final protocol validation experiment was carried out in the laboratory where 5 µL of 100pg/µL (*T. b. brucei* AnTat 1.1, *T. b. gambiense* ELIANE, *T. vivax* Y486) or 1pg/µL (*T. congolense* Forest ANR4) and 5 µL of composite West Nile *G. f. fuscipes* (containing *W. glossinidia*) DNA was spiked into the dissected tissues of insectary-reared *G. m. morsitans* (3 male and 3 female per assay target and NEC). DNA extraction was carried out following the MagnaExtract DNA extraction protocol. Multi-Tryp qPCR (dry format) was carried out on all samples, NECs and NTCs.

### Optimised field protocol and trial

#### Ethics statement

The research did not involve any human participants, domestic livestock or wild vertebrate animals. The data were produced under the auspices of the Uganda National Council of Science and Technology ethics board (“Targeting tsetse: use of targets to eliminate African sleeping sickness” Ref. Number HS 939) and a Memorandum of Understanding between the Ministry of Agriculture, Animal Industry and Fisheries (MAAIF) represented by the Coordinating Office for the Control of Trypanosomiasis in Uganda (COCTU) and the Liverpool School of Tropical Medicine (“Trypa-No! 3 – Post Elimination Tsetse Monitoring and Reactive Control, Using Tiny Targets, in Moya, Koboko, Arua, Maracha, Yumbe, Adjumani and Amuru”). In accordance with the Nagoya protocol [26], no samples were removed from the country of origin.

#### Sample Collection

Tsetse (*G. f. fuscipes*) were collected from six pyramidal monitoring traps deployed along a tributary of the Oluffe river in Maracha District, Uganda. Trap sites were situated within 30.899937° to 30.902432° North and 3.220247° to 3.221712° East. Pyramidal traps were used to regularly monitor tsetse populations at these sites for 6 years (2018 – 2024). Vector control operations, using Tiny Targets, had been implemented along part of the Oluffe river in 2012-2014, but no targets have been deployed along this part of the Olluffe since then [54]. Lying in the West Nile region of Uganda, this riverine habitat is typified by thick riverbank vegetation surrounded by mixed-use agricultural land (crop and livestock-grazing) and small villages (Fig 2). Domestic animals in the study region included cattle, goats, chickens, pigs and dogs. Flies were collected approximately three times per week for nine weeks. Collected flies were stored at 5-10°C for 20 minutes to sedate them before dissection. A stratified sampling protocol was used to select samples for further processing and analysis. In summary, where possible, equal numbers of tsetse were selected from each trap (representing equal male and female ratio) up to a maximum of 14 tsetse per day.

**Figure 2:**
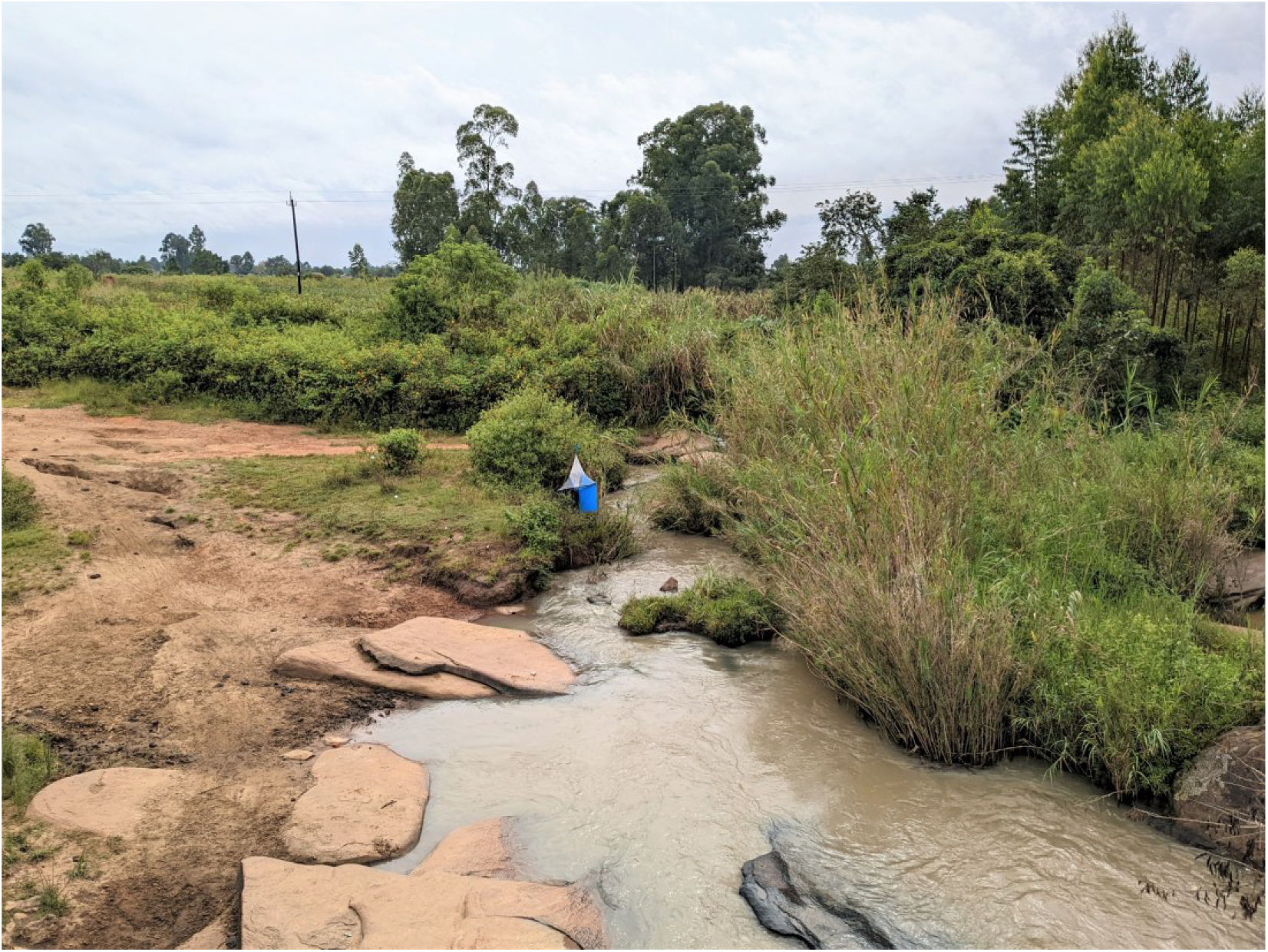
A photo showing a pyramidal trap in-situ (bottom) on the bank of the river Oluffe (trap G3-MCA). This shows the typical environment, with mixed-used agricultural land and river access point (for bathing, retrieving water and livestock drinking) on the left, and thick brush vegetation surrounding the riverbank on the right.

#### Tsetse fly dissection

To confirm infection status, all live flies were dissected and inspected by light microscopy (400X magnification) using a dark-field filter to detect trypanosome infection as described by Lloyd and Johnson [19]. Based on location of *Trypanosoma sp.* infection, flies were categorised as mouthpart (MP), midgut (MG) or salivary gland (SG) positive. Female flies were also subject to ovarian dissection and age grading, as described by Saunders [55]. The presence or absence of a visible bloodmeal (fresh or digested blood in the MG) was also recorded. Dissection equipment was cleansed with 10% bleach and rinsed in nuclease-free water between each sample. A new glass slide was used for each fly. Once dissection was complete, dissected tissue (MG, MP and SG) from each fly was placed, together, into individual collection tubes and stored at room temperature (RT) until further processing the same day (Figure 3).

**Figure 3:**
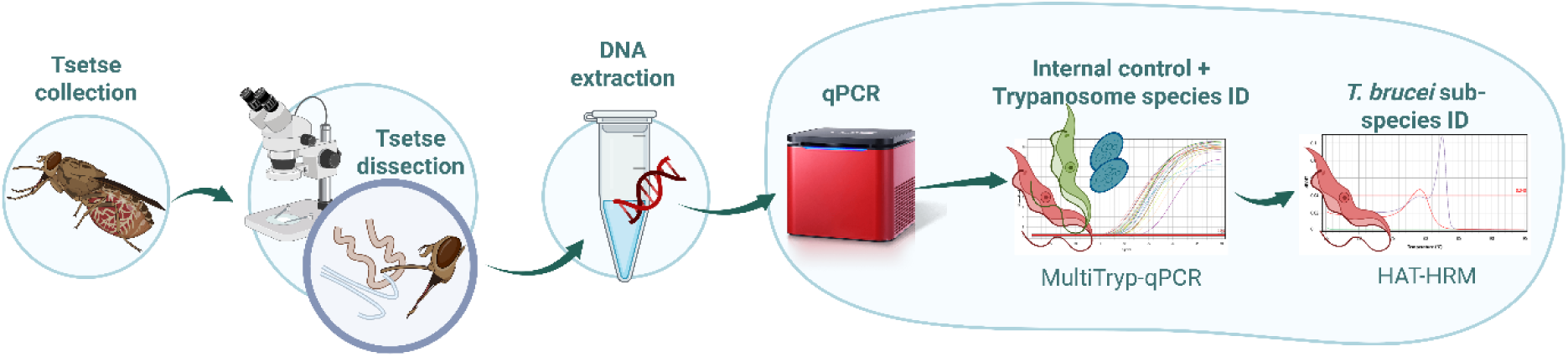
A flow diagram depicting steps in the trialled tsetse xenomonitoring protocol. Image created using biorender.com [accessed 23/04/25].

#### Field MagnaExtract DNA extraction

For field flies and negative extraction control (NEC), the optimised DNA extraction protocol used a mixture of alkaline buffers (adapted from Webster *et al* [56]), magnetic beads and wash buffers to extract and purify DNA from tsetse tissue (Figure 3). Dissected MG, MP and SGs from each fly were mixed with 70 µL of Alkaline Buffer 1 (S1 Text) and centrifuged briefly using an Appleton Mini Centrifuge (Appleton Woods, Birmingham, UK) before incubation at RT for 5 minutes. Following incubation, 130 µL of Alkaline Buffer 2 (S1 Text) was added before pulse-vortexing three times using an SLS Lab Basic Mini Vortex Mixer (SLS, Nottingham, UK). This was followed by centrifugation for 5 minutes and then further pulse-vortexing three times. A total of 50 µL supernatant was then taken forward for DNA purification. A cost-effective magnetic bead mix, adapted from a study conducted by Byrne *et al* [49] (S1 Text) was used for DNA purification. For each sample, 100 µL of this ‘SpeedBead’ solution was added and gently mixed by flushing. Following incubation at RT for 5 minutes, tubes containing sample-bead mix were placed in a magnetic rack to form a pellet of the sample. After removing supernatant, 500 µL of 70% ethanol solution was then added, the sample re-pelleted on the magnetic rack and then the supernatant removed.

Tubes were then left open to dry and remove any residual ethanol. Finally, DNA was eluted in 50 µL nuclease-free water (NFW), incubated at RT for five minutes, briefly vortexed and then centrifuged. The supernatant containing eluted DNA was then ready for subsequent analysis (Figure 3).

#### Multi-Tryp qPCR and HAT-HRM

Air-dryable qPCR reaction mixes were manufactured, dried and packaged at Liverpool School of Tropical Medicine as described previously (‘qPCR optimisation and validation’). Final optimised Multi-Tryp probe and primer concentrations were as follows: 200nM probe and 300nM primer (TBR and TCF), 200nM probe and 200nM primer (TVX) and 150nM probe and 200nM primer (UGWigg)(Table 1). Optimised HAT-HRM primer concentrations were 400nM (PLC and SRA) and 600nM (TgsGP)(Table 1).

Following transportation to PON destination, dry mixes were stored at RT out of direct sunlight until use. When required, dry mix was rehydrated following the protocol described previously (‘qPCR optimisation and validation’) and used immediately.

Thermal cycling conditions were as follows; initial denaturation at 95°C for 3 minutes followed by 40 cycles of denaturation at 95°C for 15 seconds and annealing and extension at 60°C for 35 seconds (Multi-Tryp) or 58°C for 35 seconds (HAT-HRM), followed by a high-resolution melting step between 65 – 95°C at a rate of 0.1°C (HAT-HRM only). Data were captured during the annealing and extension step. Thermocycling, fluorescence detection and data capture was carried out using a Mic and micPCR v.2.9.0 software (Bio Molecular Systems, Upper Coomera, Australia)(Figure 3).

#### Training and laboratory design

The molecular laboratory was set up in Arua, Uganda, in a small brick-built building in a residential area. The building comprised two adjacent rooms: one dedicated to DNA extraction (pre-amplification) and the other for performing qPCR (post-amplification).

Available power supplies included mains electricity, albeit subject to frequent power-cuts, and a small petrol generator. Temperature and humidity would regularly exceed 30°C and 60% respectively inside the laboratory.

At the start of the study, a two-week training workshop was carried out to equip three entomology technicians (experienced in tsetse dissection) with the knowledge and practical skills to carry out the xenomonitoring protocol independently. The first week focused on basic concepts and training in the protocol techniques, whilst the second week consisted of the technicians carrying out the full protocol and data collection under supervision. A small portable projector (Apeman M4, Apeman, UK) was used to deliver seminars on the basic concepts of DNA, molecular biology and molecular diagnostics such as targeted DNA amplification (Figure 4A). Practical training was then given in molecular techniques relevant to the protocol, in addition to quality control, research integrity and health and safety practices (Figure 4). Training was also given on the qPCR machine and software use (micPCR, BioMolecular Systems, Australia) and results interpretation, recording and reporting (Fig 4D). Standard operating procedures and laboratory design were developed in collaboration with the entomology technicians.

**Figure 4:**
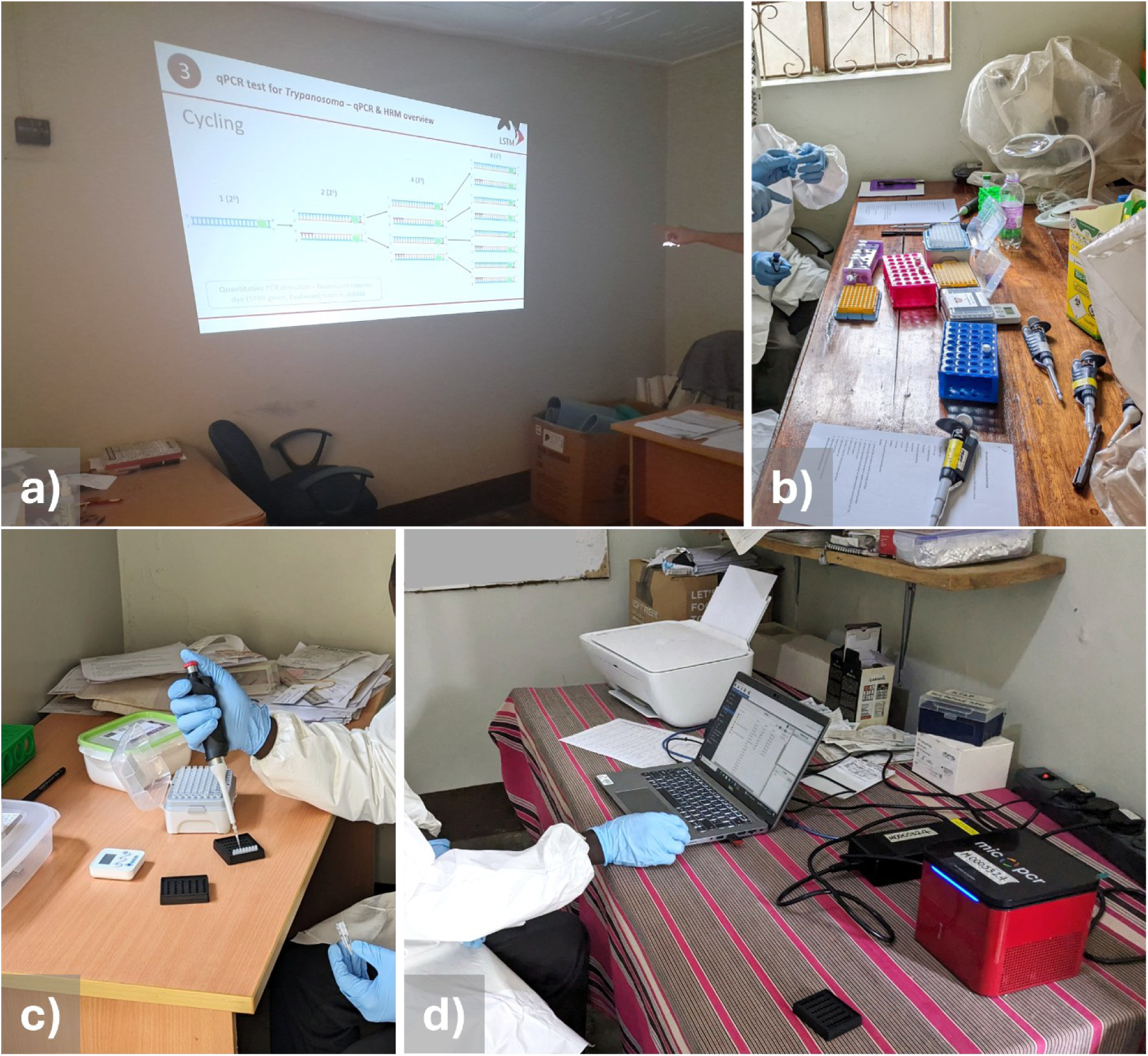
Photographs demonstrating different stages of the training workshop. Image (a) shows a lecture being given on qPCR; (b) shows training being given in DNA extraction; (c) shows the rehydration of air-dried qPCR mix (PCR Biosystems Ltd, London, UK); (d) shows a technician operating the Mic qPCR machine (BioMolecular Systems, Upper Coomera, Australia).

#### Statistical analyses

All data were collated into a centralised database in Excel (Microsoft). Further analyses and data visualisation were performed using GraphPad Prism v10. All data are presented as the mean ± standard error (SE). Spearman’s rank correlation (2-tailed) was used to determine whether there was an association between time since reagent manufacture (days) and Multi-Trip qPCR UGWigg Cq values. Pearson’s chi-square test was used to determine whether there were associations between tsetse sex, bloodfed status, trap site and detection of *Trypanosoma sp.* DNA. The Kruskal-Wallis test was used to test whether there was statistically-significant association between *Trypanosoma* detection rates and female tsetse age (ovarian category).

## Results

### Results of protocol development

#### DNA extraction optimisation

DNA extraction optimisation revealed that alkaline extraction (TM3) was the optimal method (S1 Table), with the lowest mean UGWigg-qPCR Cq and SE (Cq 21.23 ± 0.51 SE) across nine biological replicates. This was comparable to the gold standard (mean Cq 18.15 ± 0.48 SE).

#### Multi-Tryp qPCR optimisation

The optimal Multi-Tryp qPCR annealing temperature was calculated as 60°C, producing highest sensitivity across all *Trypanosoma sp.* targets (Figure 5). Optimal probe and primer concentrations were as follows: 200nM probe and 300nM primer (TBR and TCF), 200nM probe and 200nM primer (TVX) and 150nM probe and 200nM primer (UGWigg) (Figure 5). These represented the lowest primer/probes concentrations that did not significantly impact assay sensitivity. Analytical sensitivity testing using a dilution series (in spiked composite *G. f. fuscipes* DNA) showed minimal loss in assay sensitivity and specificity between wet and dried format Multi-Tryp qPCR reactions (Figure 5). Multi-Tryp qPCR 95% limit-of-detection (LOD) was calculated as 0.02 genome equivalents (1fg) per µL for TBR, 0.1 genome equivalents (10fg) per µL for TCF and 2 genome equivalents (100fg) per µL for TVX (S2 Table). The multiplex assay was also shown to be highly specific across all targets (S3 Table). Critically, no cross-reaction with *G. f. fuscipes* DNA was recorded.

**Figure 5:**
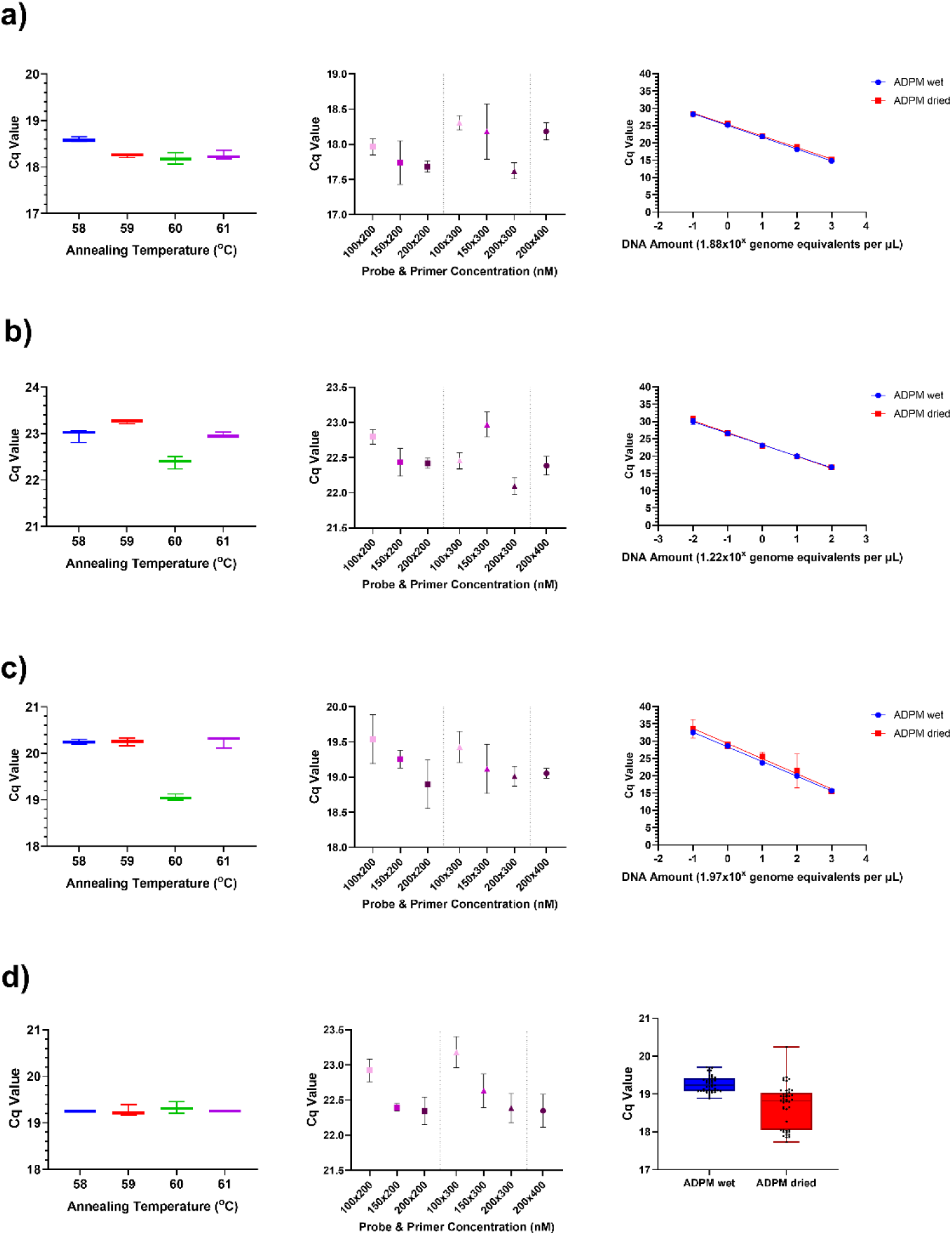
A figure displaying the mean Cq value results of optimisation experiments for targets TBR (A), TCF (B), TVX (C) and UGWigg (D) in the Multi-Tryp multiplex probe-based qPCR assay. Annealing temperature optimisation (left) and primer and probe concentration optimisation (centre) experiments were carried out using 10pg/µL *T. brucei* AnTat 1.1 DNA, 100fg/µL *T. congolense* Forest ANR4 DNA, 10pg/µL *T. vivax* DNA or a 1 in 10 dilution of composite *G. f. fuscipes* (containing *W. glossinidia*) DNA in triplicate. A five-fold log dilution series (right) of *Trypanosoma sp.* DNA spiked into composite *G. f. fuscipes* DNA was carried out using optimised wet (blue) and dry (red) Multi-Tryp air-dryable probe mix (ADPM) in triplicate. A box and whisker plot (bottom right) shows Cq value results of UGWigg-qPCR target across all spiked composite *G. f. fuscipes* samples (n=15) screened as part of the dilution series. Error bars represent min and max values.

#### HAT-HRM qPCR optimisation

The optimal HAT-HRM qPCR annealing temperature was calculated at 58°C with optimal primer concentrations at 400nM (PLC and SRA) and 600nM (TgsGP) (S2 Figure). The 95% LOD was calculated based on detection of product Tm matching expected target value: 82.5-83.5°C for PLC, 88.5-89.5°C for SRA and 91-92°C for TgsGP. Whilst sensitivity was maintained between wet and dry formats for PLC target (10pg/µL *T. brucei* DNA LOD), there was a ten-fold loss in sensitivity after drying for SRA (1 to 10pg/µL *T. b. rhodesiense* DNA LOD) and TgsGP (10 to 100pg/µL *T. b. gambiense* DNA LOD) targets in HAT-HRM (S3 Figure). Assay efficiency was difficult to calculate (S3 Figure) due to the presence of suspected primer dimers, leading to a non-specific amplicon of Tm 80.52°C (± 0.15 SE) in all reactions.

#### qPCR final validation

A final protocol validation experiment demonstrated the successful extraction and amplification of target DNA across all four assay targets (Figure 6). Multi-Tryp qPCR was shown to be sensitive and specific across TBR, TCF and UGWigg targets (Figure 6). For TVX, non-specific amplification was detected in 1/6 tsetse spiked with *T. b. gambiense* DNA (Figure 6). The reason for this is unclear, however a false-positive can be easily ruled out by cross-referencing the respective TBR result.

**Figure 6:**
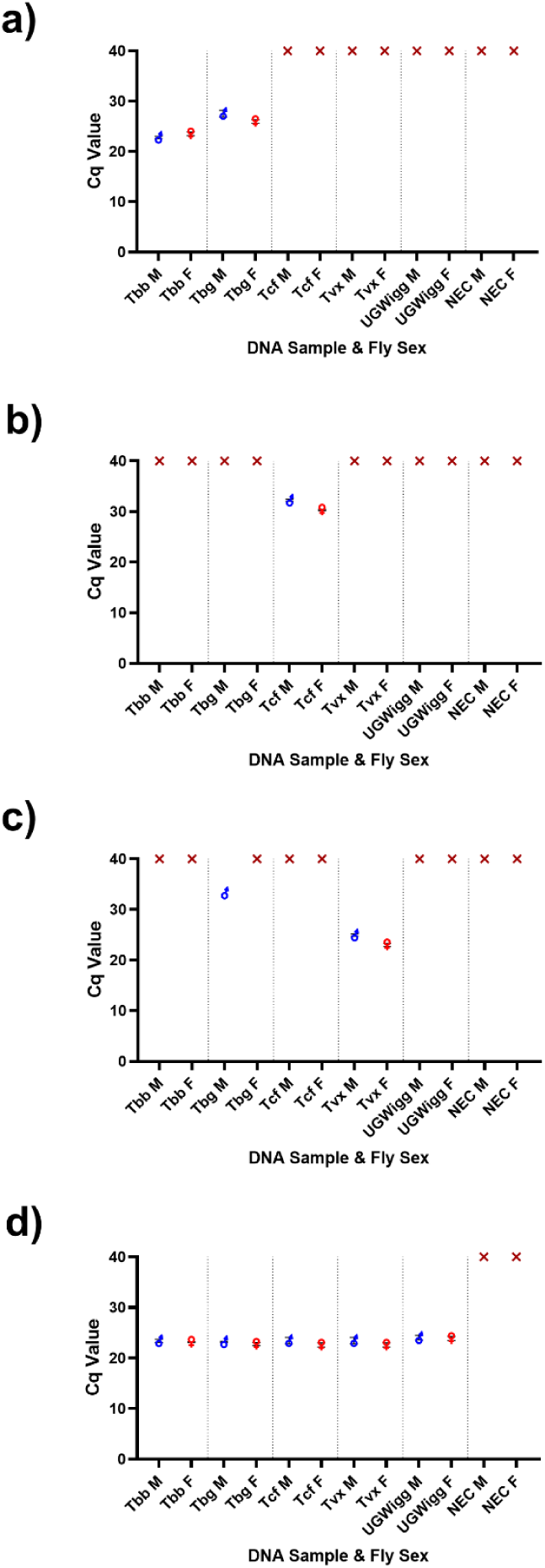
A figure displaying the Cq value results of a validation experiment for Multi-Tryp multiplex assay targets (a) TBR (*Trypanosoma brucei* s-l), (b) TCF (*T. congolense* Forest), (c) TVX (*T. vivax*) and (d) UGWigg. (*W. glossinidia*). For the experiment, 5 µL of 100pg/µL (*T. b. brucei* (Tbb) AnTat 1.1, *T. b. gambiense* (Tbg) ELIANE, *T. vivax* (Tvx) Y486), or 1pg/µL (*T. congolense* Forest (Tcf) ANR4) and 5 µL of composite West Nile *G. f. fuscipes* (containing *W. glossinidia* (UGWigg)) DNA was spiked into the dissected tissues of insectary-reared *G. m. morsitans* (3 male (M) and 3 female (F) per assay target and negative extraction control (NEC)). DNA extraction was carried out following the MagnaExtract DNA extraction protocol. All samples were then screened using the optimised dry-format Multi-Tryp qPCR assay.

**Figure 7:**
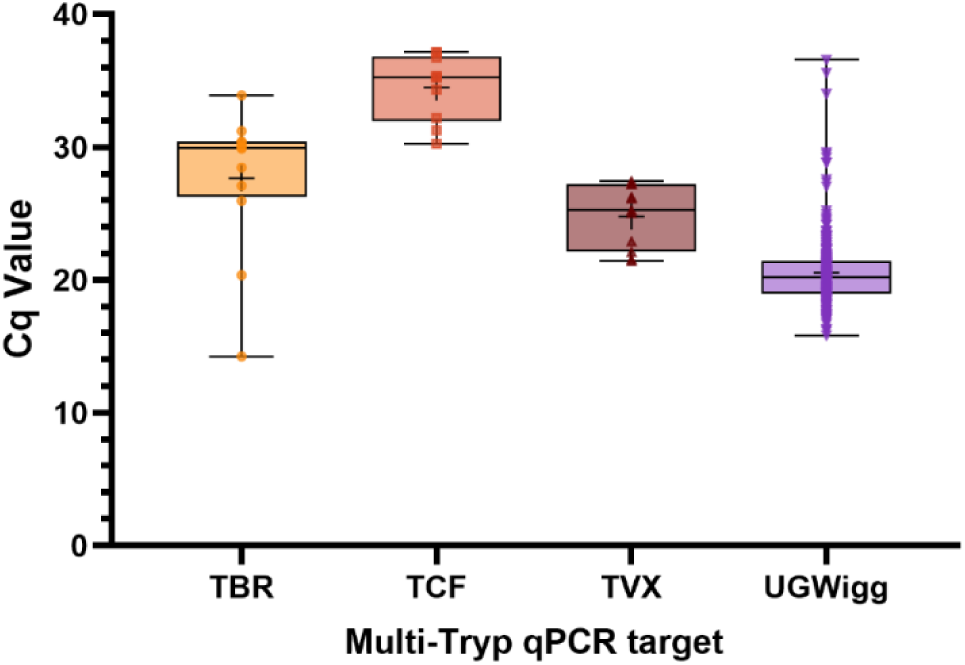
Box-and-whisker plots displaying total Cq value results of Multi-Tryp qPCR screening of wild-caught *G. f. fuscipes* (n=280) across assay targets TBR (*Trypanosoma brucei* s-l), TCF (*T. congolense* Forest), TVX (*T. vivax*) and UGWigg (*W. glossinidia*). Error bars represent minimum and maximum values, the central horizontal bar represents and median and the central cross (+) within the box represents the mean. Plotted symbols (TBR, yellow circle; TCF, orange square; TVX, brown upward triangle; UGWigg, purple downward triangle) represented individual sample Cq values.

### Field-based validation

#### DNA extraction

A total of 286 live *G. f. fuscipes* were collected for DNA extraction over nine sampling weeks between June and September 2024 with an average of nine flies processed per day. *W. glossinidia* DNA was detected in 98% (n=280) of samples, indicating high success rate of DNA extraction (Table 2). Of samples recording UGWigg amplification (n=280), the mean Cq value was 20.54 ± 0.162 SE (Figure 6). There was no significant correlation between mean UGWigg Cq value and time elapsed (days) since reagent (DNA extraction and ADPM) manufacture (*r_s_*=-0.06518, *p*=0.7322). Indicating a lack of reagent degradation over the course of the study.

**Table 2:** Multi-Tryp qPCR results (positive) from 284 tsetse across the four assay targets (*Wigglesworthia*, *T. b. brucei*, *T. congolense* Forest and *T. vivax*) against tsetse sex and blood-fed status. . CIs = confidence intervals.

| Multi-Tryp qPCR | Sex |  | Blood-fed |  | Microscopy Results |  |  | Total | Total proportion |
| --- | --- | --- | --- | --- | --- | --- | --- | --- | --- |
| Target | Male | Female | Yes | No | MG+ | SG+ | MP+ | frequency | [95% CIs] |
| UGWigg+ | 92 | 188 | 46 | 234 | 0 | 0 | 3 | 280 | 98% |
|  | 33% | 67% | 16% | 84% | 0% | 0% | 1% |  | [95.49 – 99.23] |
| TBR+ | 3 | 9 | 7 | 5 | 0 | 0 | 0 | 12 | 4.3% |
|  | 25% | 75% | 58% | 42% | 0% | 0% | 0% |  | [2.23 – 7.37] |
| TCF+ | 3 | 7 | 3 | 7 | 0 | 0 | 0 | 10 | 3.6% |
|  | 30% | 70% | 30% | 70% | 0% | 0% | 0% |  | [1.73 – 6.47] |
| TVX+ | 1 | 10 | 3 | 8 | 0 | 0 | 3 | 11 | 3.9% |
|  | 9% | 91% | 27% | 73% | 0% | 0% | 27% |  | [1.98 – 6.92] |
| Total frequency | 92 | 192 | 47 | 237 | 0 | 0 | 3 |  |  |
| Total proportion | 32% | 68% | 17% | 83% | 0% | 0% | 1% |  |  |

### Detection of *Trypanosoma sp.* DNA

DNA of *Trypanosoma sp.* causing AAT were also detected successfully. Across all samples amplifying UGWigg DNA (n=284), 10.7% (n=30) contained *Trypanosoma sp.* DNA, of which 4.3% (n=12) contained *T. brucei* DNA, 3.6% (n=10) *T. congolense* Forest and 3.9% (n=11) *T. vivax* (Table 2). Of these, three flies were found to contain DNA of two *Trypanosoma sp.*; *T. brucei* and *T. congolense* Forest (n=1), *T. brucei* and *T. vivax* (n=1) and *T. congolense* Forest and *T. vivax* (n=1). HAT-HRM screening (S4 Figure) revealed that 67% (n=8/12) *T. brucei* s-l samples were likely to be *T. b. brucei* (mean Tm 82.78°C ± 0.098°C). A further four (of 12) *T. brucei* s-l samples failed to amplify the single copy GPI-PLC gene, indicating that insufficient DNA was present for accurate sub-species identification. No *T. b. gambiense* or *T. b. rhodesiense* DNA was detected in any samples.

Dissection and microscopy results revealed no salivary gland or midgut infections, however all mouthpart infections (n=3) also tested positive for *T. vivax* DNA. There was a statistically significant association between blood-fed tsetse and detection of *Trypanosoma sp.* DNA in tsetse (χ_2_=13.597, *p*<0.001). Compared to the proportion of blood-fed tsetse across the total population (17%, n=46), there were higher proportions of blood-fed tsetse (27-58%) in samples also containing *Trypanosoma sp.* DNA (Table 2).

Of 57 negative extraction controls (NECs) across 33 Multi-Tryp qPCR runs, 5.26% (n=3) recorded UGWigg amplification (mean Cq 32.38 ± 2.39 SE). No amplification was recorded in NECs across *Trypanosoma* targets (TBR, TCF and TVX). No amplification was recorded in any Multi-Tryp qPCR NTCs (n=33). The lack of contamination highlights both the aptitude of the technicians and the robustness of the protocol, despite the basic nature of the laboratory.

## Discussion

### Overview

This study demonstrates the successful development, optimisation and implementation of a tsetse molecular xenomonitoring protocol in a low-resource point-of-need setting. Utilising novel components, such as a low-cost alkali and magnetic bead-based DNA extraction method, air-dryable probe-based qPCR reactions and a small, portable qPCR machine ensured that molecular xenomonitoring could be undertaken in a remote location without reliance on cold-chain or large high-powered equipment. The qPCR results show the sensitive and consistent detection of trypanosome DNA over the course of the study, with no evidence of carry-over contamination at either the DNA extraction or qPCR stages. The high success rate of DNA extraction (98%) highlights the robustness of the optimised protocol and the consistency and diligence of the technicians in following standard operating procedures. Despite having no prior experience in molecular biology, the technicians were fully independent after two weeks of training and adapted quickly to new protocol methods. The high protocol competency shown after just one week of training is evidence that the workshop could be condensed to a one-week intensive course, to be brought in line with the WHO gHAT diagnostic test target product profile [28].

### *Trypanosoma sp.* detection and epidemiological considerations

There was no *T. b. gambiense* DNA detected in the tsetse screened, further evidencing the elimination of gHAT in this area [4,7,57]. This is consistent with other livestock and tsetse xenomonitoring studies previously carried out in the area which also did not detect *T. b. gambiense* [7–10,13]. Nonetheless, the reduced sensitivity performance of the dried-format HAT-HRM assay must be kept in mind (S3 Figure). The optimised *T. b. gambiense* LOD of dried-format HAT-HRM (100pg/µL, or 2x10^6^ genome equivalents per mL) was 100 times less sensitive than the previously-reported LOD (10^4^ tryps per mL) that used an HRM-specific master mix (Thermo-start ABgene, Rochester, New York, USA) [39]. The 10-fold higher sensitivity of the PLC target compared to TgsGP target also introduces the possibility of low-concentration *T. b. gambiense* DNA being mis-identified as *T. b. brucei*. The low sensitivity of molecular tests for *T. b. gambiense* is a recognised problem [17] and remains a major obstacle in gHAT xenomonitoring. The fact that the Multi-Tryp probe-based assay did not record loss in sensitivity when dried, in contrast to the HRM assay, suggests that the HAT-HRM assay may benefit from redevelopment to a multiplex probe-based assay for this particular air-dryable qPCR mix.

Although no *T. b. gambiense* DNA was detected, the study has demonstrated that there are several different livestock pathogenic trypanosomes circulating in the area (Table 2). The *Trypanosoma sp.* detection prevalences of 3.6% - 4.3% ([1.73% - 7.73%] 95% CI; or 10.7% in total) (Table 2) are higher than those reported for West Nile *G. f. fuscipes* previously.

Cunningham *et al* have previously found *T. brucei* s-l detection prevalences of 0.1 – 1.8%, *T. congolense* prevalences of 0.3 – 2.7% and *T. vivax* prevalences of 1 – 2.4%. The high sensitivity of the Multi-Tryp qPCR assay is a likely explanation for these differences. Previous studies used ITS-PCR (in combination with TBR-PCR or RIME-LAMP) to detect *Trypanosoma* [7,11,13], which targets a lower copy-number region than the highly sensitive minichromosome satellite repeat targets of TBR, TCF and TVX [58]. High sensitivity assays can lead to problems in differentiating infected and uninfected tsetse [24], yet this sensitivity can also be viewed as an advantage; detecting trypanosome DNA in blood-fed or sub-infectious tsetse allows researchers to gain insights into which parasites are circulating in local host populations, also known as ‘xenosurveillance’ [59]. This is evidenced in the current study by the statistically significant association found between blood-fed tsetse and detection of *Trypanosoma* DNA (χ2=13.597, *p*<0.001), in addition to the generally low quantity of target DNA (mean Cq values 24.78 – 34.49; Figure 6) detected in the samples. This suggests that the qPCR assay may be detecting *Trypanosoma* in the bloodmeals rather than maturely infected flies. Whilst xenosurveillance does not allow for accurate quantifiable estimates of infection prevalence, it can be used to explore trypanosome species richness and diversity in a given area.

The consistent and frequent detection of AAT pathogen DNA is a finding of concern for local livestock keepers and indicates the presence of infected animals in the area. Our findings are consistent with previous AAT surveillance studies in West Nile region, which also detected *T. brucei* s-l, *T. congolense* and *T. vivax* in domestic animals and livestock [6–9]. A 2014 census revealed that 83.1% of households across Maracha district engaged in livestock farming, with two thirds of total households relying on subsistence farming as the main source of income [60]. AAT is therefore a potential livelihood and food security risk for local communities. Future screening of tsetse for bloodmeal sources, domestic animals for trypanosomes and local surveys on trypanocide and insecticide use could further assess the risk and impact of AAT in the region.

### Performance and feasibility of the molecular xenomonitoring protocol

Although probe-based chemistry is more expensive, it enabled the multiplexing of four sensitive and specific assays with easy-to-interpret amplification results (Figure 5, Figure 6). Cost was instead minimised by optimising Multi-Tryp qPCR for a low (10 µL) final reaction volume, meaning that approximate cost of screening per sample was approximately £0.40 (∼US$0.53). The air-dryable qPCR mix performed consistently given the challenging ambient temperature (>30°C) and humidity (>60%) conditions. No association was found between time since manufacture and UGWigg-Cq value (*r_s_*=-0.06518, *p*=0.7322) and assays continued to amplify *Trypanosoma* and *W. glossinidia* DNA at 92 days (13 weeks) post-manufacture. A previous study testing air-dryable qPCR assay sensitivity over a 12-week period reported degradation from eight weeks in one of two assays stored at RT [61]. Whilst beyond the scope of the current study, the Multi-Tryp qPCR would benefit from a similar study testing assay integrity over time to determine maximum shelf-life.

This study also describes the successful development and use of *W. glossinidia* as a tsetse qPCR endogenous control target. *W. glossinidia* is an obligate endosymbiont of *Glossina sp.*, transmitted to progeny through maternal milk secretions [62]. It is the dominant endosymbiont, accounting for an estimated >99% of the tsetse microbiome [63] and is found in the midgut where *Trypanosoma* also reside. These qualities make it an attractive choice for an internal qPCR control. For this use we have assumed that *W. glossinidia* is present in all tsetse and at relatively equal and constant quantity throughout adult life stages. However, there are known differences in bacterial load between tsetse species. Previous studies analysing the microbiota of Ugandan tsetse found that *G. f. fuscipes* collected from Murchison Falls national park had 10X higher density of *W. glossinidia* than *G. m. morsitans* or *G. pallidipes* collection from the same region [63]. There was also some variation in the *W. glossinidia* density between *G. f. fuscipes* individuals (n=6), however this remained within 10^5^ – 10^6^ genomic equivalents [63]. The use of *W. glossinidia* as an internal control in future xenomonitoring studies, particularly those in other regions or involving different tsetse species, will require careful results interpretation.

In addition to the 98% DNA extraction success rate evidenced by sample UGWigg amplification, the lack of significant change in UGWigg Cq values over time is also indicative of the stability of DNA extraction reagents. Whilst the DNA extraction method had to be completed in several steps, it required low-powered equipment only (mini centrifuge and mini vortex mixer) and the approximate cost was just £0.10 (∼US$0.13) per sample, which is considerably less expensive than commercial kits. The total time-to-result (from field collection to qPCR result) was approximately three to four hours, depending on the number of samples being processed. These aspects would likely change if the protocol was redeveloped to process whole tsetse, without the need for prior dissection. Previous studies determined that mechanical lysis and/or overnight incubation was required to extract DNA from whole tsetse using the MagnaExtract method [10]. An alternative solution would be the use of isothermal amplification such as LAMP or RPA. Isothermal amplification methods are the obvious ‘field-friendly’ assay choice, requiring few resources, having higher tolerance to inhibitors, capable of screening crude ‘extraction-free’ sample preparations and yet able to produce high sensitivity and specificity results [11,64,65].

However, these assays can be challenging to multiplex and do not allow the quantification of target DNA, which can be a critical aspect of differentiating mature infected and uninfected tsetse [24]. There are benefits and advantages to both isothermal and qPCR methods, with final selection dependent on user needs and study objectives.

Another option to streamline sample processing would be a sample pooling strategy. Pooling has been trialled successfully in tsetse previously using LAMP assays [11,66] and can reduce processing time and DNA extraction cost whilst allowing for high-throughput analyses. Pooling capability largely depends on assay sensitivity. Whilst the RIME-LAMP assay is more sensitive than Multi-Tryp qPCR, with a reported LOD of 0.1 *T. brucei* per Ml in tsetse [11] compared to 19 *T. brucei* genome equivalents per mL (S2 Table), pooling is worth exploring alongside further DNA extraction optimisation.

Whilst this PON lab is not as mobile as other ‘lab in a bag’ studies [36,67,68], we have demonstrated reliable long-term use over nine weeks. Although the laboratory had access to mains electricity, power cuts were frequent which necessitated the use of a small mobile generator during qPCR machine use (which required at least one and half hours of uninterrupted supply per run). The micPCR machine is designed to run on battery power, which was not trialled in the current study but would provide an even more sustainable and mobile solution. Whilst a fridge was used in this study to chill tsetse prior to dissection, this step was not strictly necessary and no other reagents or equipment required cold-chain storage. A major problem facing all PON diagnostic test platforms in sub-Saharan Africa is the inconsistency or lack of supply chains for consumables and resources. Linking to this, are the difficulties in repairing and servicing equipment without in-country technical support. These are issues recognised in the WHO *T. b. gambiense* diagnostic test TPP [28]. In order for molecular diagnostics to remain sustainable in the long-term without the need for external re-supply, supply chain improvement needs to be considered as part of disease surveillance capacity strengthening.

Molecular xenomonitoring at the PON is a growing field. With the cost of molecular technology decreasing, there are hopes that access to these methods can continue to grow. In this study we have demonstrated the sensitive and consistent tsetse xenomonitoring in a low-resource field station over several weeks at an approximate cost of £0.50 (∼US$0.66) per sample. As ever, the challenge in xenomonitoring remains the low parasite prevalence necessitating large sample sizes and need for low-cost high-throughput techniques. Future research should focus on sample pooling strategies and continuing to develop more sensitive assays for the detection of *T. b. gambiense*. To ensure the long-term sustainability of PON molecular xenomonitoring, further work is also required to strengthen equipment and consumables supply chains, support training initiatives and further explore the use of alternative power sources.

## Data availability statement

All data generated during this project is available at https://figshare.com/s/9487d03e84be9a47b376.

## Supporting information captions

**S1 Text: Protocols for the making of Alkaline Buffers and SpeedBead Mix for the purposes of DNA extraction.** Protocols for Alkaline Buffers 1 and 2 was adapted from Webster *et al* [56]. Protocol for SpeedBead Mix was adapted from Byrne *et al* [49].

## Acknowledgements

I would like to extend my gratitude to Irene, Angelo, David, Joshua, Tiger and all other members of the extended Arua team who facilitated in the smooth running of this project. Thank you also to Prof Liam Morrison, Prof Annette Macleod, Prof Wendy Gibson and Prof Andrew Jackson for providing *Trypanosoma sp.* DNA samples.

---

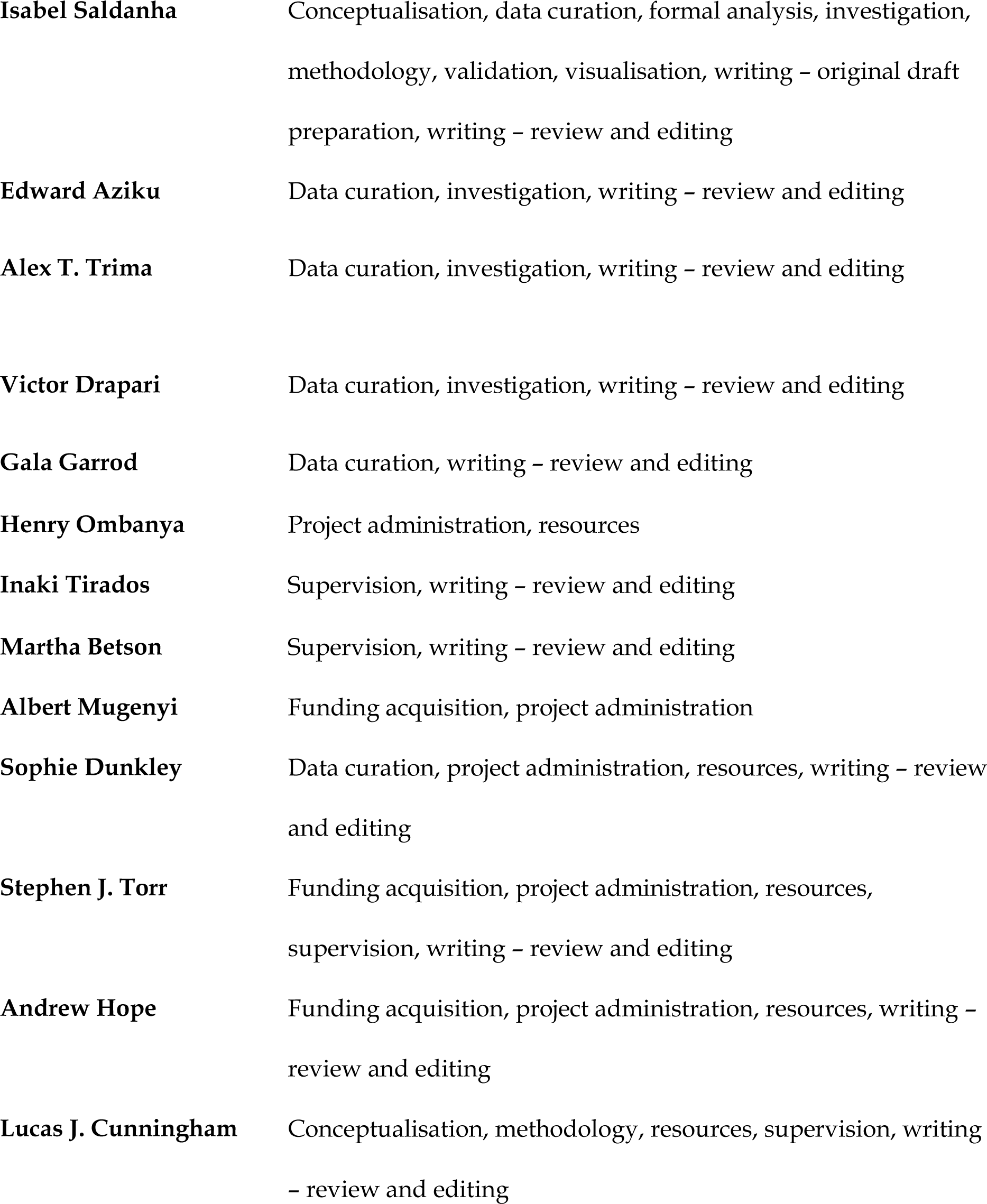

**S1 Figure:** Protocol for DNA extraction optimisation and evaluation experiments, testing three experimental methods (TM 1, TM 2 and TM 3) against a gold standard (DNeasy Blood and Tissue kit; QIAGEN, Hilden, Germany). RT = room temperature.

**S2 Figure:** **Charts plotting the mean Cq value results (i) and product melt temperatures (Tm) (ii) of primer concentration optimisation experiments for HAT-HRM assay PLC (a), SRA (b) and TgsGP (c) targets, tested in triplicate**. DNA screened was *T. b. brucei*, *T. b*. *gambiense* and *T. b. rhodesiense* at 10pg/µL. (d) displays results of HAT-HRM betaine optimisation experiments in pure *T. b. gambiense* (Tbg) DNA (100pg/µL), Tbg DNA spiked into *G. f. fuscipes* composite DNA (10pg/µL Tbg) and pure *G. f. fuscipes* composite DNA. Error bars represent standard error, horizontal bars represent the mean.

**S3 Figure:** Plots displaying analytical sensitivity results for wet format (i), dried format (ii) and wet versus dry format (iii) optimised HAT-HRM assays across five serial log dilutions (0.01 - 100 pg/µL) of triplicate target *Trypanosoma sp.* DNA: (a) *T. b. brucei* (PLC), (b) *T. b. rhodesiense* (PLC + SRA) and (c) *T. b. gambiense* (PLC + TgsGP). In charts (i) and (ii), melt temperatures (Tm) for each amplified target product (PLC, blue square; SRA, red circle; TgsGP, green triangle) are plotted. Horizontal bars represent the mean, red shaded value on x-axis represents the 95% limit-of detection. In charts (iii), HAT-HRM mean Cq values and standard curves are plotted for wet (blue, circle symbol) and dry (red, square symbol) format reactions. Error bars represent standard error.

**S4 Figure:** Images showing amplification traces for UGWigg (A) and TBR (B) targets as part of Multi-Tryp qPCR screening of eight samples, two negative extraction controls and one negative template control (NTC). All samples (n=8) amplified *Wigglesworthia* target DNA (A) and one sample tested positive for *T. brucei* s-l DNA (B). The melt profile resulting from HAT-HRM screening (C) of this sample purple trace) and NTC (red trace) showed that the sample is likely to be *T. b. brucei* based on a single melt peak at 82.89°C.

**S1 Table:** A table displaying Cq values obtained from *Wigglesworthia*-qPCR (S1 Figure) screening of nine insectary-reared *G. m. morsitans* DNA extracted using three field extraction methods (TM1-3), compared against a gold standard extraction method (QIAGEN DNeasy Blood and Tissue; S1 Figure). TM1 = TE buffer, TM2 = lysis buffer and TM3 = alkaline extraction. SE = standard error.

**S2 Table:** Tables displaying analytical sensitivity limit-of-detection (LOD) results for the optimised dry-format Multi-Tryp qPCR against three trypanosome targets *T. brucei* s-l (a), *T. congolense* Forest (b) and *T. vivax* (c). SD = standard deviation, PP = proportion positive.

**S3 Table:** Tables displaying optimised dry-format Multi-Tryp qPCR analytical specificity testing results across the four targets; TBR - *T. brucei* s-l (a), TCF-*T. congolense* Forest (b), TVX - *T. vivax* (c) and UGWigg – *W. glossinidia* (d).

