## Supplementary S1 Figure for "Development and pilot application of a point-of-need molecular xenomonitoring protocol for tsetse (*Glossina sp.*) in a low-resource setting"

**TM 1: TE Buffer**

- Place dissected tsetse tissues in 1.5 mL microcentrifuge tube and briefly centrifuge
- Add 200 µl of TE buffer
- Incubate at RT for 5 minutes

**TM 2: Lysis Buffer**

- Place dissected tsetse tissues in 1.5 mL microcentrifuge tube and briefly centrifuge
- Add 100 µl of Cell Lysis Solution (Merck Millipore, Burlington US)
- Incubate at RT for 5 minutes
- Add 100 µl of TE buffer

**TM 3: Alkaline extraction**

- Place dissected tsetse tissues in 1.5 mL microcentrifuge tube and briefly centrifuge
- Add 70 µL of alkaline buffer 1 (S1), to the sample
- Incubate at RT for 5 minutes
- Add 130 µL of alkaline buffer 2 (S1), to the sample
- Pulse vortex 3 times
- Incubate at RT for 10 minutes

**DNA purification using SpeedBead mix (S2)**

- Pulse-vortex 10 times
- Centrifuge at 8,000rpm for 5 minutes
- Pipette out 50 µl of supernatant to new tube
- Add 100 µl of SpeedBeads, flush to mix.
- Incubate at RT for 5 mins
- Pellet with magnetic rack.
- Remove supernatant
- Add 500 µl of 70% ethanol solution, flush to mix.
- Pellet with magnetic rack.
- Remove supernatant.
- Allow beads to dry 5-20 mins
    1. NOTE: beads should appear matt, not glossy
- Elute DNA by adding 50ul of nuclease free water to dry beads
- Allow to incubate at RT for 5 mins, vortex and spin
- Use 2 µl of extract in 12 µl qPCR

***Glossina morsitans morsitans*-specific *Wigglesworthia glossinidia* qPCR**

A qPCR assay targeting the tsetse obligate endosybiont *W. glossinidia* was carried out to assess the quality and quantity of DNA extraction. The primers were designed against a *G. m. morsitans*-specific *W. glossinidia* entry (accession number CP003315.1) targeting the 16s rRNA gene region. Primers were designed using Primer3. For the qPCR reactions, 2 µL of eluted DNA was mixed with 6 µL 2X Brilliant III Ultra-Fast qPCR master mix (Agilent Technologies, Santa Clara, US) 0.4 µM forward (WIG_GMM_F: 5'-CTTAGTGCATTATTGTTTTATCCTTTAGATTTAGG-3') and reverse (WIG_GMM_R: 5'-TGTGGCGTCCCCTAGGGGGTTCGA-3') primers and nuclease-free water added to a 12 µL final reaction volume. Thermal cycling conditions were as follows; initial denaturation at $95^{o}$C for 3 minutes followed by 40 cycles of denaturation at $95^{o}$C for 10 seconds and annealing and extension at $59^{o}$C for 12 seconds.

**S1 Figure:** Protocol for DNA extraction optimisation and evaluation experiments, testing three experimental methods (TM 1, TM 2 and TM 3) against a gold standard (DNeasy Blood and Tissue kit; QIAGEN, Hilden, Germany). RT = room temperature.
