## Supplementary S1 Table for "Development and pilot application of a point-of-need molecular xenomonitoring protocol for tsetse (*Glossina sp.*) in a low-resource setting"

| Replicate number | Test method | | | Gold standard |
|---|---|---|---|---|
| | **TM1** | **TM2** | **TM3** | **Qiagen** |
| **1** | 28.52 | | 20.11 | 17.52 |
| **2** | **-** | 23.95 | 21.66 | 17.50 |
| **3** | **-** | 21.30 | 19.16 | 18.30 |
| **4** | **-** | 21.71 | 23.16 | 18.77 |
| **5** | 37.02 | 21.65 | 20.68 | 18.29 |
| **6** | **-** | 23.12 | 19.11 | 17.39 |
| **7** | 29.52 | 21.57 | 21.01 | 21.63 |
| **8** | **-** | 27.51 | 22.15 | 17.15 |
| **9** | **-** | 21.98 | 23.10 | 16.76 |
| **Mean Cq** | **31.69** | **22.85** | **21.13** | **18.15** |
| **Cq SE** | **2.68** | **0.74** | **0.51** | **0.48** |

**S1 Table: A table displaying Cq values obtained from *Wigglesworthia*-qPCR (S1 Figure) screening of nine insectary-reared *G. m. morsitans* DNA extracted using three field extraction methods (TM1-3), compared against a gold standard extraction method (QIAGEN DNeasy Blood and Tissue; S1 Figure).** TM1 = TE buffer, TM2 = lysis buffer and TM3 = alkaline extraction. SE = standard error.
