## Supplementary S1 Text for "Development and pilot application of a point-of-need molecular xenomonitoring protocol for tsetse (*Glossina sp.*) in a low-resource setting"

**Alkaline Buffer 1:** 0.1M NaOH, 0.3mM EDTA, pH13.0
NaOH, molar mass: 39.997 g/mol.
EDTA, molar mass: 292.2438 g/mol.
To make 10X stock solution:
Dissolve 4g NaOH in 80mL distilled water.
Add 0.876g EDTA.
Add distilled water to 100mL.
Dilute 10X stock solution 1:10 for working buffer. Ensure that final pH is 13.

**Alkaline Buffer 2:** 0.1M Tris-HCL, pH 7.0
Tris-HCL, molar mass: 121.14 g/mol
To make 10X solution:
Dissolve 12.1g in 80mL distilled water.
Use 5M NaOH to make solution up to 100mL and increase pH (if necessary).
Make 0.1M working solution by diluting 10X stock 1:10 with distilled water. Ensure that final pH is 7.

**SpeedBead Mix Protocol**

1. Transfer 1ml of Sera-Mag Carboxylate-Modified Magnetic Beads and Speedbeads (Cytiva, Marlborough, US) to a 1.5ml microcentrifuge tube.
2. Place microcentrifuge tubes on a magnetic rack until beads are separated.
3. Remove supernatant.
4. Add 1ml TE (pH 7.5-8.0) to beads, remove from magnetic rack, mix by pipetting up and down and then return to magnetic rack.
5. Remove supernatant.
6. Repeat steps 4 & 5.
7. Add 1ml TE (pH 7.5-8.0) to beads, remove from magnetic rack, mix by pipetting up and down, but DO NOT return to magnet.
8. Add 9g PEG-8000 to a new 50ml conical tube.
9. Add 2.92g NaCl to conical.
10. Add 500ul 1M Tris-HCl to conical.
11. Add 100ul 0.5M EDTA to conical.
12. Fill conical to 49ml using ddH20.
13. Mix conical for 5 minutes until PEG goes into solution.
14. Add 25ul Tween 20 to conical and mix.
15. Add SpeedBeads and TE solution from step 7 to conical and mix.

Fill conical to 50ml with ddH20.

**S1 Text: Protocols for the making of Alkaline Buffers and SpeedBead Mix for the purposes of DNA extraction.** Protocols for Alkaline Buffers 1 and 2 was adapted from Webster *et al* [56]. Protocol for SpeedBead Mix was adapted from Byrne *et al* [49].
