## Supplementary figures and images for "Development and pilot application of a point-of-need molecular xenomonitoring protocol for tsetse (*Glossina sp.*) in a low-resource setting"

### Supplementary S2 Figure

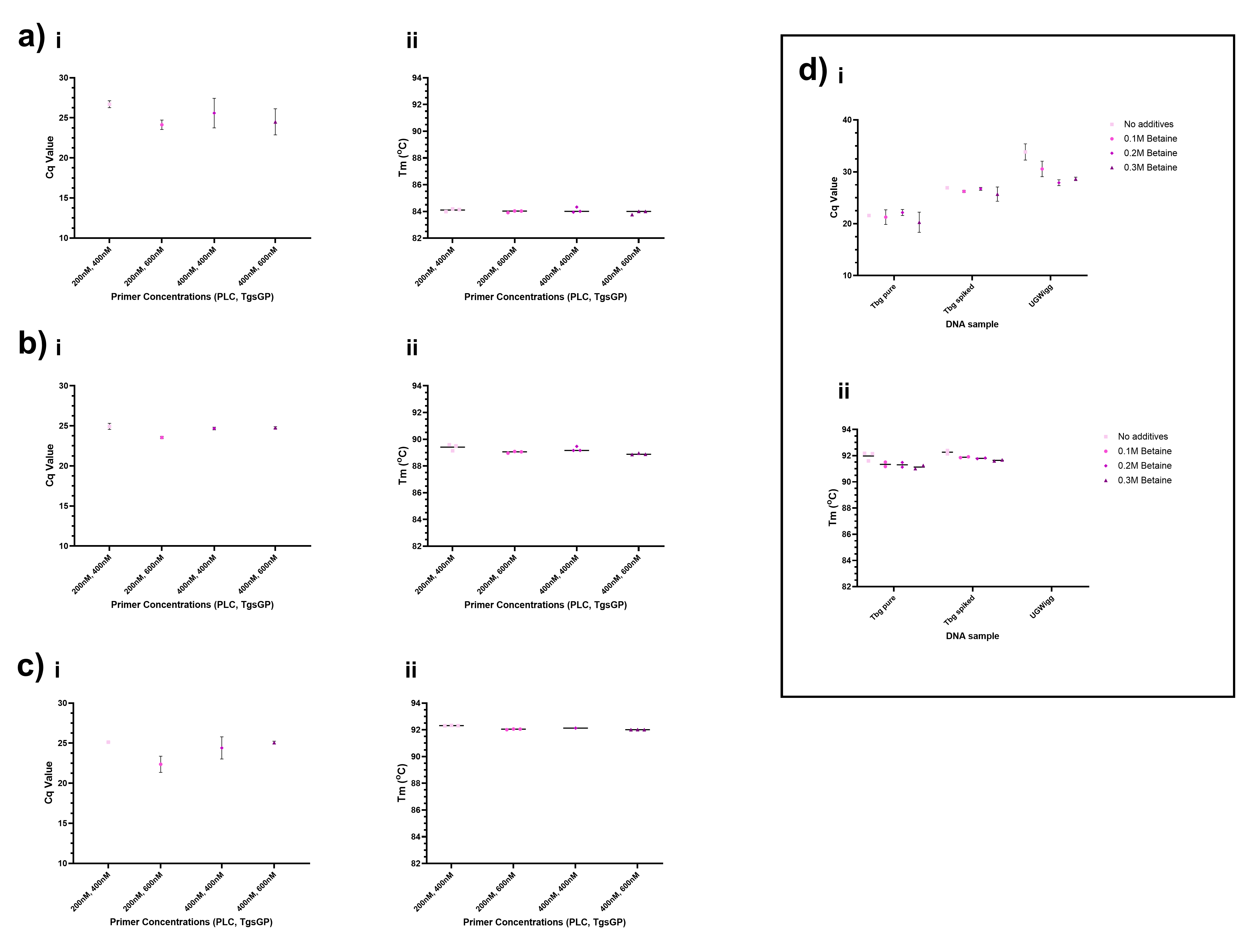

### Supplementary S3 Figure

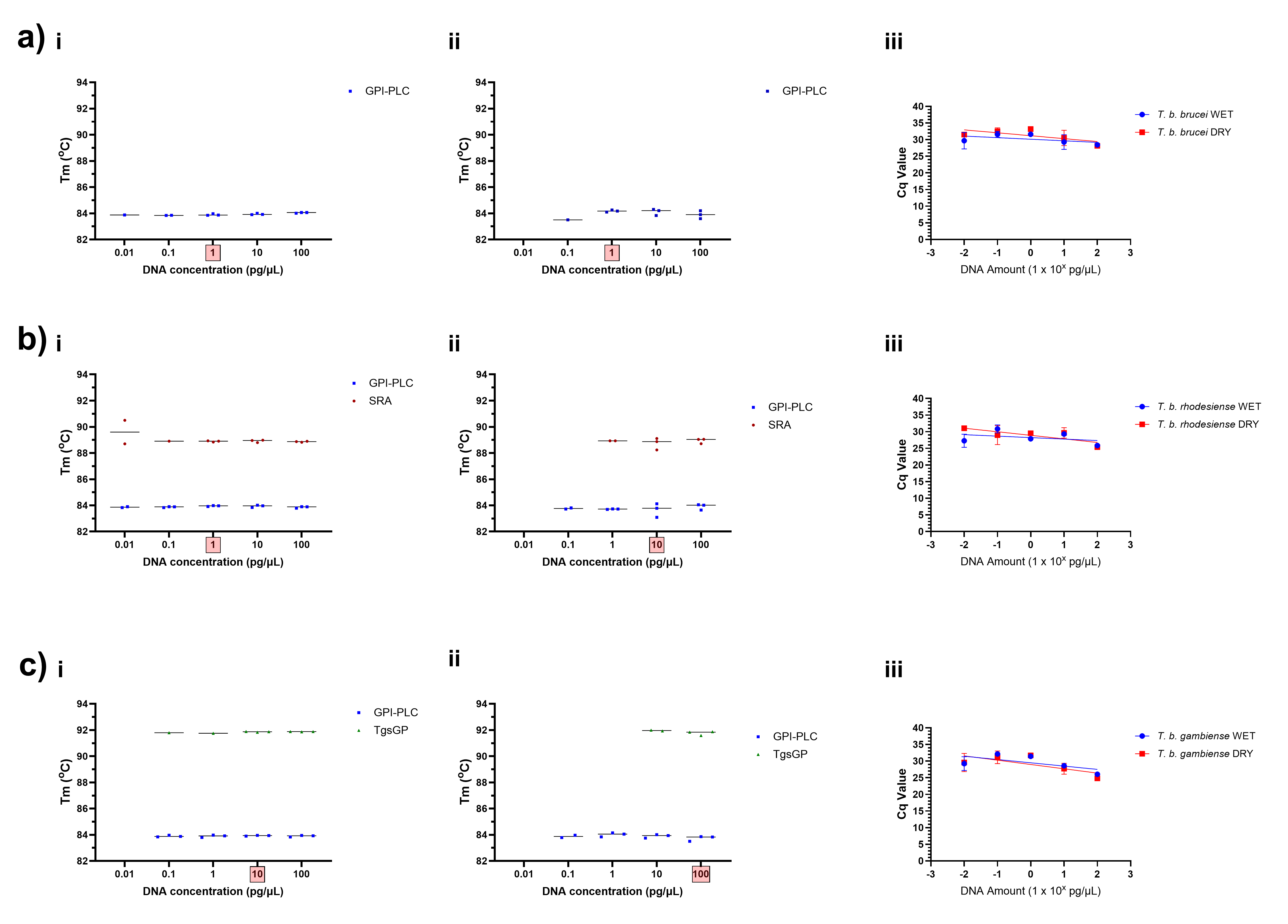
