## Supplementary S2 Table for "Development and pilot application of a point-of-need molecular xenomonitoring protocol for tsetse (*Glossina sp.*) in a low-resource setting"

**a)** *T. brucei* (TBR)

| DNA conc (copies/µL) | $1.88x10^{0}$ | $1.88x10^{-1}$ | $1.88x10^{-2}$ | $1.88x10^{-3}$ | $1.88x10^{-4}$ |
|---|---|---|---|---|---|
| **Mean Cq** | 26.59 | 29.78 | 35.40 | NA | NA |
| **SD** | 1.40 | 0.18 | 2.22 | NA | NA |
| **Amplification** | 4 | 4 | 4 | 0 | 0 |
| **PP (%)** | 100% | 100% | 100% | 0% | 0% |

**b)** *T. congolense* Forest (TCF)

| DNA conc (copies/µL) | $1.22x10^{0}$ | $1.22x10^{-1}$ | $1.22x10^{-2}$ | $1.22x10^{-3}$ | $1.22x10^{-4}$ |
|---|---|---|---|---|---|
| **Mean Cq** | 29.04 | 31.40 | 34.86 | NA | NA |
| **SD** | 0.33 | 1.66 | 1.84 | NA | NA |
| **Amplification** | 4 | 4 | 2 | 0 | 0 |
| **PP (%)** | 100% | 100% | 50% | 0% | 0% |

**c)** *T. vivax* (TVX)

| DNA conc (copies/µL) | $1.97x10^{1}$ | $1.97x10^{0}$ | $1.97x10^{-1}$ | $1.97x10^{-2}$ | $1.97x10^{-3}$ |
|---|---|---|---|---|---|
| **Mean Cq** | 25.08 | 27.87 | 31.64 | NA | NA |
| **SD** | 1.06 | 0.31 | 1.01 | NA | NA |
| **Amplification (/4)** | 4 | 4 | 2 | 0 | 0 |
| **PP (%)** | 100% | 100% | 50% | 0% | 0% |

**S2 Table: Tables displaying analytical sensitivity limit-of-detection (LOD) results for the optimised dry-format Multi-Tryp qPCR against three trypanosome targets *T. brucei* s-l (a), *T. congolense* Forest (b) and *T. vivax* (c).** SD = standard deviation, PP = proportion positive.
