## Supplementary S3 Table for "Development and pilot application of a point-of-need molecular xenomonitoring protocol for tsetse (*Glossina sp.*) in a low-resource setting"

a)

| Species | Sub-species | Strain | DNA conc. | Target (Y/N) | Amplification (total = 3) | Mean Cq. |
|---|---|---|---|---|---|---|
| ***T. brucei*** | *brucei* | AnTat 1.1 | 1ng/µL | Y | 3 | 11.81 |
| ***T. brucei*** | *gambiense* | ELIANE | 1ng/µL | Y | 3 | 17.45 |
| ***T. brucei*** | *rhodesiense* | Z3 | 1ng/µL | Y | 3 | 10.72 |
| ***T. congolense*** | Savannah | IL3000 | 1ng/µL | N | 0 | - |
| ***T. congolense*** | Forest | ANR3 | 10fg/µL | N | 0 | - |
| ***T. congolense*** | Kilifi | WG84 | 1ng/µL | N | 0 | - |
| ***T. simiae*** | | TV008 | 1ng/µL | N | 0 | - |
| ***T. simiae*** | Tsavo | 114 | 1ng/µL | N | 0 | - |
| ***T. vivax*** | | Y486 | 1ng/uL | N | 0 | - |
| ***G. f. fuscipes* (West Nile) + *W. glossinidia* composite** | | | Unknown | N | 0 | - |

b)

| Species | Sub-species | Strain | DNA conc. | Target (Y/N) | Amplification (total = 3) | Mean Cq. |
|---|---|---|---|---|---|---|
| ***T. brucei*** | *brucei* | AnTat 1.1 | 1ng/µL | N | 0 | - |
| ***T. brucei*** | *gambiense* | ELIANE | 1ng/µL | N | 0 | - |
| ***T. brucei*** | *rhodesiense* | Z3 | 1ng/µL | N | 0 | - |
| ***T. congolense*** | Savannah | IL3000 | 1ng/µL | N | 0 | - |
| ***T. congolense*** | Forest | ANR3 | 10fg/µL | Y | 3 | 25.71 |
| ***T. congolense*** | Kilifi | WG84 | 1ng/µL | N | 0 | - |
| ***T. simiae*** | | TV008 | 1ng/µL | N | 1 | 38.27 |
| ***T. simiae*** | Tsavo | 114 | 1ng/µL | N | 0 | - |
| ***T. vivax*** | | Y486 | 1ng/uL | N | 0 | - |
| ***G. f. fuscipes* (West Nile) + *W. glossinidia* composite** | | | Unknown | N | 0 | - |

c)

| Species | Sub-species | Strain | DNA conc. | Target (Y/N) | Amplification (total = 3) | Mean Cq. |
|---|---|---|---|---|---|---|
| ***T. brucei*** | *brucei* | AnTat 1.1 | 1ng/µL | N | 0 | - |
| ***T. brucei*** | *gambiense* | ELIANE | 1ng/µL | N | 0 | - |
| ***T. brucei*** | *rhodesiense* | Z3 | 1ng/µL | N | 0 | - |
| ***T. congolense*** | Savannah | IL3000 | 1ng/µL | N | 0 | - |
| ***T. congolense*** | Forest | ANR3 | 10fg/µL | N | 0 | - |
| ***T. congolense*** | Kilifi | WG84 | 1ng/µL | N | 0 | - |
| ***T. simiae*** | | TV008 | 1ng/µL | N | 0 | - |
| ***T. simiae*** | Tsavo | 114 | 1ng/µL | N | 0 | - |
| ***T. vivax*** | | Y486 | 1ng/uL | Y | 3 | 12.56 |
| ***G. f. fuscipes* (West Nile) + *W. glossinidia* composite** | | | Unknown | N | 0 | - |

d)

| Species | Sub-species | Strain | DNA conc. | Target (Y/N) | Amplification (total = 3) | Mean Cq. |
|---|---|---|---|---|---|---|
| ***T. brucei*** | *brucei* | AnTat 1.1 | 1ng/µL | N | 0 | - |
| ***T. brucei*** | *gambiense* | ELIANE | 1ng/µL | N | 0 | - |
| ***T. brucei*** | *rhodesiense* | Z3 | 1ng/µL | N | 0 | - |
| ***T. congolense*** | Savannah | IL3000 | 1ng/µL | N | 0 | - |
| ***T. congolense*** | Forest | ANR3 | 10fg/µL | N | 0 | - |
| ***T. congolense*** | Kilifi | WG84 | 1ng/µL | N | 0 | - |
| ***T. simiae*** | | TV008 | 1ng/µL | N | 0 | - |
| ***T. simiae*** | Tsavo | 114 | 1ng/µL | N | 0 | - |
| ***T. vivax*** | | Y486 | 1ng/uL | N | 0 | - |
| ***G. f. fuscipes* (West Nile) + *W. glossinidia* composite** | | | Unknown | Y | 3 | 19.27 |

**S3 Table: Tables displaying optimised dry-format Multi-Tryp qPCR analytical specificity testing results across the four targets; TBR - *T. brucei* s-l (a), TCF- *T. congolense* Forest (b), TVX - *T. vivax* (c) and UGWigg – *W. glossinidia* (d).**
