## Supplementary S4 Figure for "Development and pilot application of a point-of-need molecular xenomonitoring protocol for tsetse (*Glossina sp.*) in a low-resource setting"

**a)** *Wigglesworthia glossinidia*

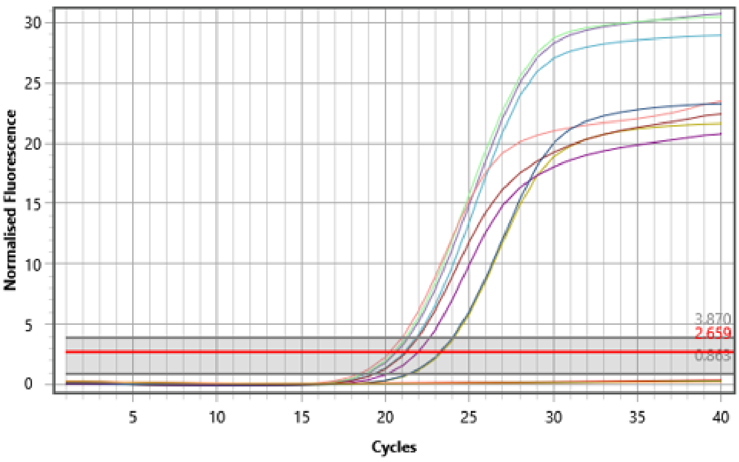


**b)** *T. brucei* s-l

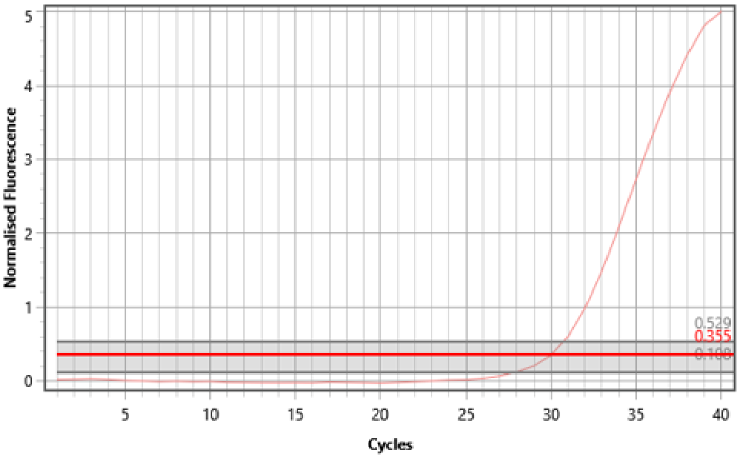


**c)** HAT-HRM

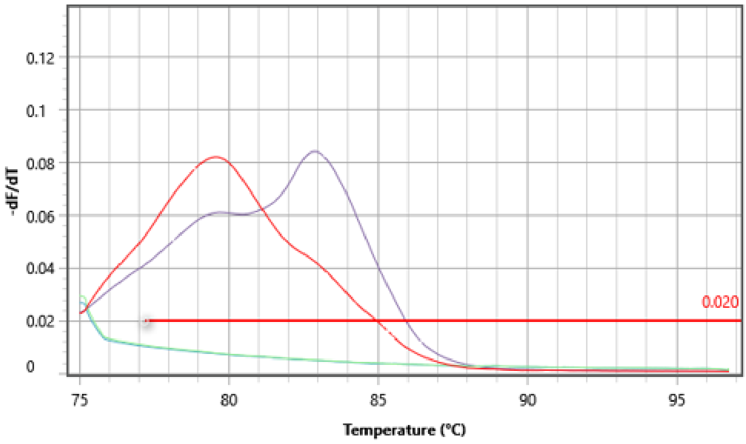


**S4 Figure: Images showing amplification traces for UGWigg (A) and TBR (B) targets as part of Multi-Tryp qPCR screening of eight samples, two negative extraction controls and one negative template control (NTC).** All samples (n=8) amplified *Wigglesworthia* target DNA (A) and one sample tested positive for *T. brucei* s-l DNA (B). The melt profile resulting from HAT-HRM screening (C) of this sample purple trace) and NTC (red trace) showed that the sample is likely to be *T. b. brucei* based on a single melt peak at 82.89$^{o}$C.
